# Why scaling up uncertain predictions to higher levels of organisation will underestimate change

**DOI:** 10.1101/2020.05.26.117200

**Authors:** James A. Orr, Jeremy J. Piggott, Andrew Jackson, Jean-François Arnoldi

## Abstract

Uncertainty is an irreducible part of predictive science, causing us to over- or underestimate the magnitude of change that a system of interest will face. In a reductionist approach, we may use predictions at the level of individual system components (e.g. species biomass), and combine them to generate predictions for system-level properties (e.g. ecosystem function). Here we show that this process of scaling up uncertain predictions to higher levels of organization has a surprising consequence: it will systematically underestimate the magnitude of system-level change, an effect whose significance grows with the system’s dimensionality. This stems from a geometrical observation: in high dimensions there are more ways to be more different, than ways to be more similar. This general remark applies to any complex system. Here we will focus on ecosystems thus, on ecosystem-level predictions generated from the combination of predictions at the species-level. In this setting, the ecosystem’s dimensionality is a measure of its diversity. We explain why dimensional effects do not play out when predicting change of a single linear aggregate property (e.g. total biomass), yet are revealed when predicting change of non-linear properties (e.g. absolute biomass change, stability or diversity), and when several properties are considered at once to describe the ecosystem, as in multi-functional ecology. Our findings highlight and describe the counter-intuitive effects of scaling up uncertain predictions, effects that will occur in any field of science where a reductionist approach is used to generate predictions.

## 1 Introduction

In natural sciences, uncertainty of any given prediction is ubiquitous (Dovers and Handmer, 1992). When considering predictions of change, uncertainty has directional consequences: uncertain predictions will lead to either over- or underestimation of actual change. The reductionist approach to complex systems is to gather and use knowledge about individual components before scaling up predictions to the system-level (Levins and Lewontin, 1985; Wu et al., 2006). Although scaling up to higher levels of organisation is general to the study of any complex systems, it is particularly well-defined in ecology. In this field, knowledge about the components at lower levels of organisation (individuals, populations) is commonly used to understand the systems at higher levels of organisation (communities, ecosystems) (Loreau, 2010; Woodward et al., 2010).

An unbiased prediction of an individual component is one that makes no systematic bias towards over- or underestimation for that component (Box 1). But what happens when we scale up unbiased predictions to higher levels of organisation? If we do not systematically underestimate the change of individual components, will this still be true when considering many components at once? When addressing this question, one must be wary of basic intuitions as the problem is inherently multi-dimensional, thus hard to properly visualize.

As a thought experiment, consider two ecological communities, one species-poor (low dimension) and the other species-rich (high-dimension). Both communities experience perturbations that change species biomass, and we assume that we have an unbiased prediction for this change, up to some level of uncertainty. We then scale up our predictions to the community-level, focusing on the change in Shannon’s diversity index, caused by the perturbations. By comparing predicted and observed change we can quantify the degree of underestimation of our predictions, at the species and community-level. If we simulate this thought experiment (Fig. 1 and Appendix S4) we observe the following puzzling results, which motivate our subsequent analysis. Predictions of species biomass change may be unbiased (bottom row of Fig. 1), but when scaled up to the system level for the species-rich community, but not the species-poor community, we see a clear bias towards underestimation of change (top right corner of Fig. 1).

**Figure 1.**
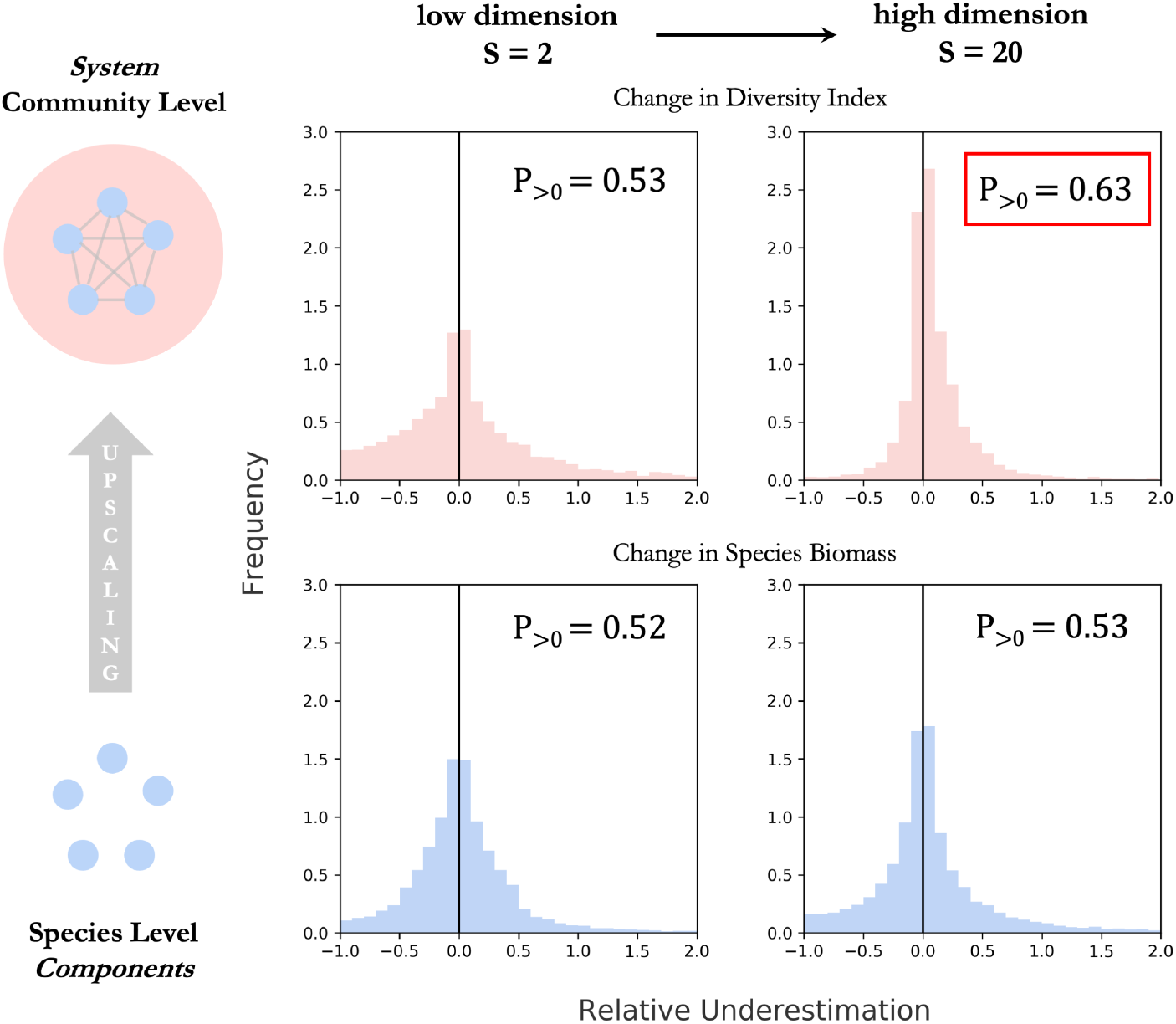
Simulated communities of 2 species (left) and 20 species (right) experienced 1000 perturbations (change in species biomass), for which we assume unbiased predictions at the species-level. Uncertainty around those predictions is simulated as a random terms of zero mean, independent across species. Histograms show the distribution of relative underestimation, defined as the difference between realized and predicted change expressed relatively to the predicted magnitude of change. By construction, there is no bias towards underestimation at the species level (bottom row). We then scale up our predictions to the community level to generate predictions for Shannon’s diversity index (top row). For the first, species poor community, this upscaling does not generate any bias. However, for the species rich community a bias emerges as 63% of realizations show an underestimated magnitude of change. In this article, we explain in depth the statistical mechanisms behind this bias.

As we shall explain in depth, the reason for this emergent bias is that *in high dimensions there are more ways to be more different, than ways to be more similar*. Our goal is to make this statement quantitative and generally relevant to ecological problems. We start from a geometric approach showing that, in two dimensions, our claim can be visualised to reveal a positive relationship between magnitude of uncertainty and underestimation of change. Visualisation is only possible in low dimensions, but a more abstract reasoning demonstrates that as dimensionality increases so does the bias towards underestimation, which is further strengthened by larger uncertainty. We note that dimensionality is not necessarily an integer value. We propose that the effective dimensionality most relevant to ecological upscaling of predictions is not the number of species, but instead is a specific diversity metric, the Inverse Participation Ratio (IPR) (Suweis et al., 2015; Wegner, 1980), comparable (but not equivalent) to Hill’s diversity indices (Hill, 1973).

We then explain why the effect of dimensionality depends on how change is measured at the system-level (Box 1). If a single linear function is used to aggregate components (e.g. total biomass), dimensionality has no effect. An unbiased prediction for individual components trivially scales up to produce an unbiased system-level prediction. But this is not true in general. Non-linear functions (e.g. Shannon’s diversity index as in Fig. 1), can remain sensitive to dimensional effects. Predictions of change of these properties, even if constructed from unbiased predictions of individual components will be systematically underestimated. The significance of this effect will depend on the relative significance of non-linearities in the function of interest.

On simulated examples we will examine the behaviour of common ecosystem-level properties: diversity, stability and total biomass. More generally, we emphasise that dimensional effects will occur as soon as system-level change is measured as a change in multiple properties at once (whether they are linear or not), as is the case in multi-functional descriptions of ecosystems (Manning et al., 2018).

As a seemingly different kind of ecological case-study, we then revisit core questions of multiple-stressor research in the light of our theory. In this field, there is a clear prediction (additivity of stressor effects), a high prevalence of uncertainty about the the way stressors interact (resulting in non-additivity) and, ultimately, great interest in the ecosystem-level consequences of non-additive stressor interactions (synergism or antagonism) (Côté et al., 2016; Jackson et al., 2016; Piggott et al., 2015). Expressed in this context, our theory predicts the generation of bias towards synergism when multiple-stressor predictions are scaled up to higher levels of organisation.

Research has primarily focused on the causes of uncertainty, working hard to reduce it (Petchey et al., 2015). Here we take a complementary approach by investigating the generic consequences of uncertainty, regardless of the nature of the system studied or the underlying causes of uncertainty. Our theory becomes more relevant as the degree of uncertainty increases, which makes it particularly relevant for ecological problems. But, in fact, our findings could inform any field of science that takes a reductionist approach in the study of complex systems (e.g. economics, energy supply, demography, finance – see Box 2), demonstrating how dimensional effects can play a critical role when scaling up predictions.

### Box 1: Lexicon of Concepts

**Reductionist view of complex systems**

- *Components:* Individuals variables *B_i_* that together form a system (e.g. biomass of *S* species and abiotic compartments forming an ecosystem).
- *System state:* Point in state *space*, represented as a vector *B* = (*B*_1_,…, *B_S_*) jointly describing all system components.
- *Difference* (*or magnitude of change*) *between states*: the Euclidean distance ||*B* − *B′* || between two joint states *B* and *B′*.

**Scaling up uncertain predictions**

- *Relative error:* Magnitude of error caused by uncertainty relative to the magnitude of predicted change.
- *Aggregate system-level property:* Scalar function of the joint state (e.g. total biomass or diversity index)
  *Linear aggregate property:* Linear function of joint state variables (e.g. total biomass).
  *Non-linear property:* Non-linear function of joint state variables (e.g. diversity index).
- *Scaled up prediction:* A prediction made for the joint state, or a scalar property of the joint state, based on individual predictions for components.
- *Unbiased prediction:* A prediction that, despite uncertainties, does not systematically overestimate or underestimate the magnitude of change (of a joint state, a system component or an aggregate property).

**Multi-functional view of complex systems**

- Multivariate description of a complex system, based on multiple aggregate properties, or *functions* (production, diversity, respiration) instead of individual components (species biomass and abiotic compartments). The state of the system is the joint state *F* = (*F*_1_,,…, *F_S_F__*) of *S_F_* functions. Difference between states is the distance between two joint functional states *F* and *F′*.

## 2 Geometric Approach

The central claim of this article is that *in high dimensions there are more ways to be more different, than ways to be more similar*. We propose an implication: *a system-level prediction based on unbiased predictions for individual components, will tend to underestimate the magnitude of system-level change*.

To understand these statements, it is useful to take a geometrical approach to represent the classic reductionist perspective, starting in two dimensions (Fig. 2a). Picture two intersecting circles in a system’s state-space (one blue, one red in Fig. 2). The first, blue circle is centred on the system’s initial state and its radius corresponds to the predicted magnitude of change. The second, red circle is centred at the predicted state (which lies on the blue circle) and its radius corresponds to the magnitude of realized error of the prediction, in other words, the realized outcome of the uncertainty associated with the prediction (red circle in Fig. 2). The actual final state is thus somewhere on that red circle. If it falls outside the blue circle, the prediction has underestimated the magnitude of change. The proportion of the red circle lying outside of the blue circle measures the proportion of possible configurations that will lead to an underestimation of change. In other words, for a given magnitude of error caused by uncertainty, this portion of the circle represents the states that are more different from the initial state than predicted. As the relative magnitude of error increases (as the red circle’s diameter becomes larger, relative to that of the blue circle) this proportion grows (Fib. 2a).

**Figure 2.**
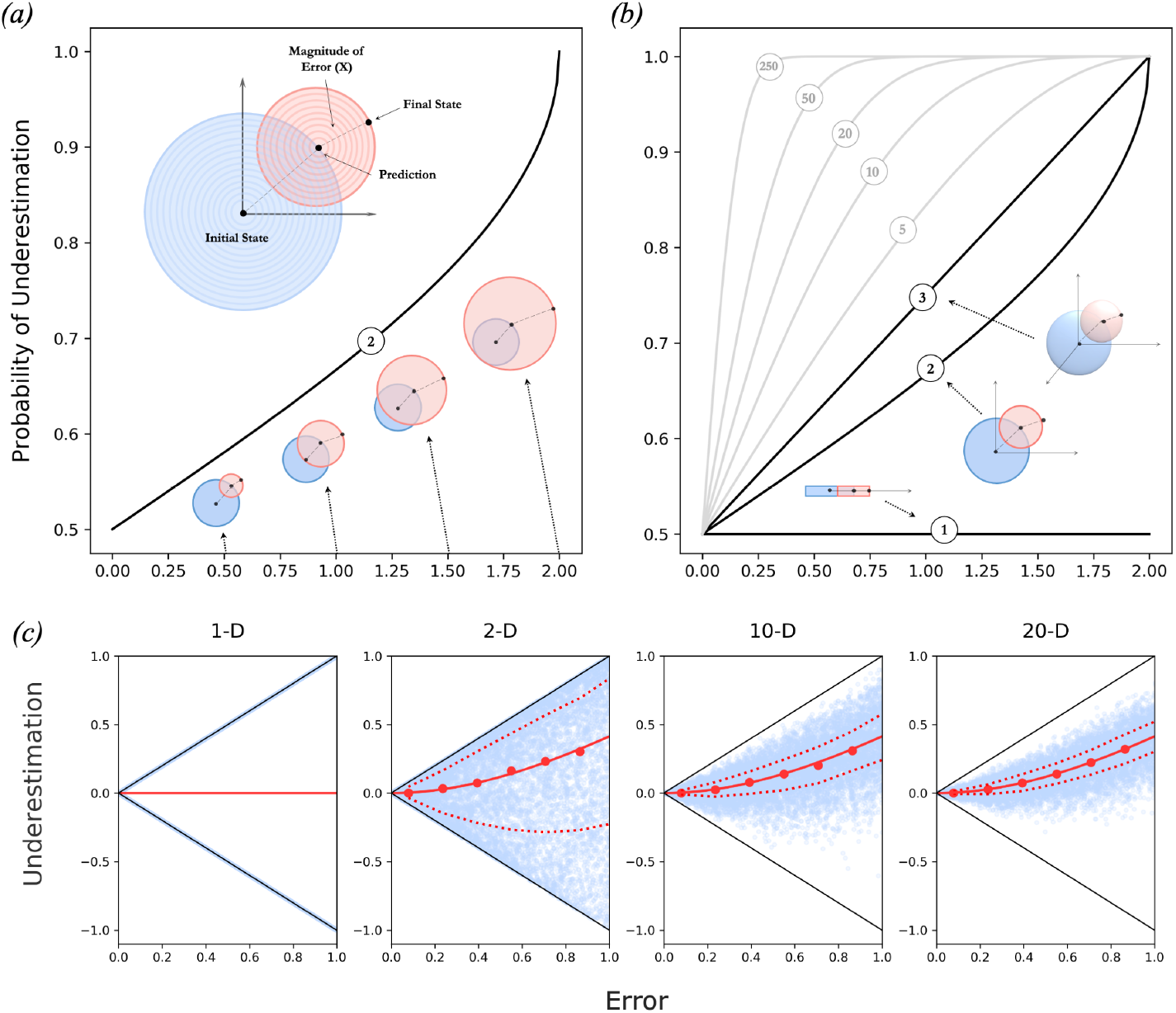
***(a)*** Already in two dimensions, the probability of underestimation increases as uncertainty increases. The centre of the blue circle is the initial state (its actual value is irrelevant) and its radius is defined by the predicted magnitude of change. The point at the centre of the red circle corresponds to the predicted state, while its radius represents the magnitude of error made by the prediction. By definition, final states thus fall on the edges of the red circle. If a final state falls inside the blue circle then there has been an overestimation of change (it is closer to the initial state than what was predicted). If a final state falls outside the blue circle (as in the figure) then there has been an underestimation of change (it farther from the initial state than what was predicted). When uncertainty is small, error will be small thus the radius of the red circle is small, and the probability of underestimation is close to 0.5. As uncertainty (thus error) increases, however, there is increasing bias towards underestimation. Eventually when error is twice as large as the prediction only underestimation is possible. ***(b)*** This relationship between uncertainty and underestimation is strengthened by dimensionality. As dimensions increases there become even more ways to be more different than ways to be more similar. Each curve corresponds to the probability of underestimation as a function of error for different dimensions labeled as circled numbers. For a fixed amount of error the probability of underestimation will increase with dimension. ***(c)*** The relationship between the relative magnitude of error (*x*) and the relative magnitude of underestimation (*y*) based on uniform sampling of 1-D, 2-D, 10-D and 20-D intersecting hyper-spheres defined by unbiased but uncertain predictions. The boundaries of this relationship are plotted in black and the median expectation 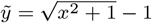 as derived from Eq. (4) is plotted in red (except for 1-D where it does not apply). Blue points are simulated results, red points are the actual median values and dashed lines show the quantiles for vertical subsets of the simulated data. As dimensionality increases the width of the distribution decreases and converges towards its median, which effectively increases the probability of underestimation (b).

In three dimensions these two intersecting circles become two intersecting spheres. The proportion of interest is the surface of the spherical cap lying outside of the sphere centred on the initial state. Here, a non-intuitive phenomenon occurs: with the same radii as in the 2D case, in 3D there are now more configurations leading to underestimation. As dimensions increase this proportion increases, until the vast majority of possible states now lie in the domain where change in underestimated (Fig. 2b). This result can be made quantitative from known expressions for the surface of hyper-spherical caps. This gives us an analytical expression for the proportion of configurations leading to an underestimation of change, as a function of the relative magnitude of error (*x*) and dimension (*S*):

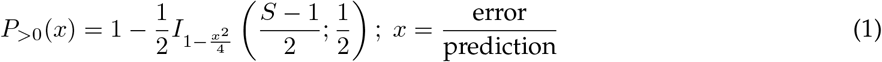

In the above equations || · || stands for the standard Euclidean norm of vectors^1^, and *I_s_* (*a, b*) is the cumulative function of the *β*-distribution (Appendix S2). This is what we mean by *in high dimensions there are more ways to be more different, than ways to be more similar*. To see how this relates to the scaling up of unbiased predictions of individual components (Box 1), we now take a statistical approach. Suppose we uniformly sample the intersecting circles, spheres and hyper-spheres defined above and drawn in Fig. 2. The proportion Eq. (1) becomes a probability, the probability of having underestimated change. This uniform sampling is precisely what happens if the uncertainty of individual variables are independent random normal variables with zero mean (a particular case of an unbiased uncertainty at the component level, see Appendix S2). This justifies our second claim: *a system-level prediction based on unbiased predictions for individual components, will tend to underestimate the magnitude of change of the system state*.

This reasoning is geometrical, and relies on a computation of the surface of classic shapes such as hyper-spheres and spherical caps. But the core mechanism behind the behaviour of the probability of underestimation is more general and in a sense, simpler. To see that, let us take a step back and analyse the relative magnitude of underestimation, defined as:

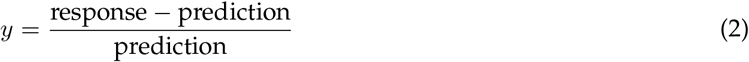

Given an angle *θ* between prediction and error vectors (resp. the vectors that point from initial to predicted state, and from predicted state to realized state) we can rearrange Eq. (2) as:

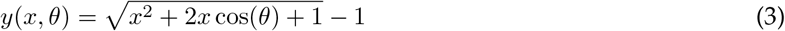

the term cos*θ* can take any values between −1 and +1. For the sake of simplicity, in what follows we will suppose that its mean and median are zero. This is the case if the errors associated with individual variables are drawn from independent symmetric distributions centred on zero (unbiased and unskewed predictions at the component level). In this case the median relationship between error (*x*) and underestimation (*y*) is:

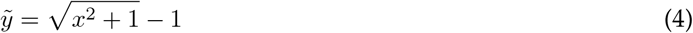

which is strictly positive as soon the error *x* is non zero. This holds true in all dimensions greater than one, which can be seen in Fig. 2c. The median underestimation 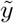 does not depend on dimension, but the probability of underestimation, *P*(*y* > 0; *x*), does. Indeed, *P*(*y* > 0; *x*) is driven by the distribution of the random term cos*θ* in Eq. (3). If this distribution is narrow, realisations of *y* will fall close to 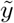. Because the latter is positive and increases predictably with *x*, so will the probability of any realised *y* to be positive. A known fact from random geometry is that, given a random isotropic vector (i.e. a vector whose direction is uniformly distributed on the sphere), its angle *θ* with any other given vector satisfies

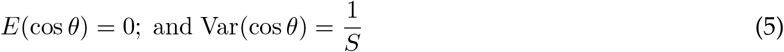

In other words, in high dimensions random vectors are approximately orthogonal, up to a variance inversely proportional to the dimension of state-space. In our context, this corresponds to normal i.i.d. distributions of errors, a particular case of independent unbiased and unskewed predictions. This explains why the probability of underestimation increases in Fig. 3b with both dimension *S* and error *x*. In what follows we use the expression for the variance in Eq. (5) as a *definition* of *effective dimension*. In doing so, we can free ourselves from the strict Euclidean representation of Fig. 2, and generalize the theory beyond i.i.d. normal error distributions. This will be useful when applying our theory to ecological problems, where components are the biomass of species, are their contribution to ecosystem change are not equivalent, thus errors not i.i.d.

**Figure 3.**
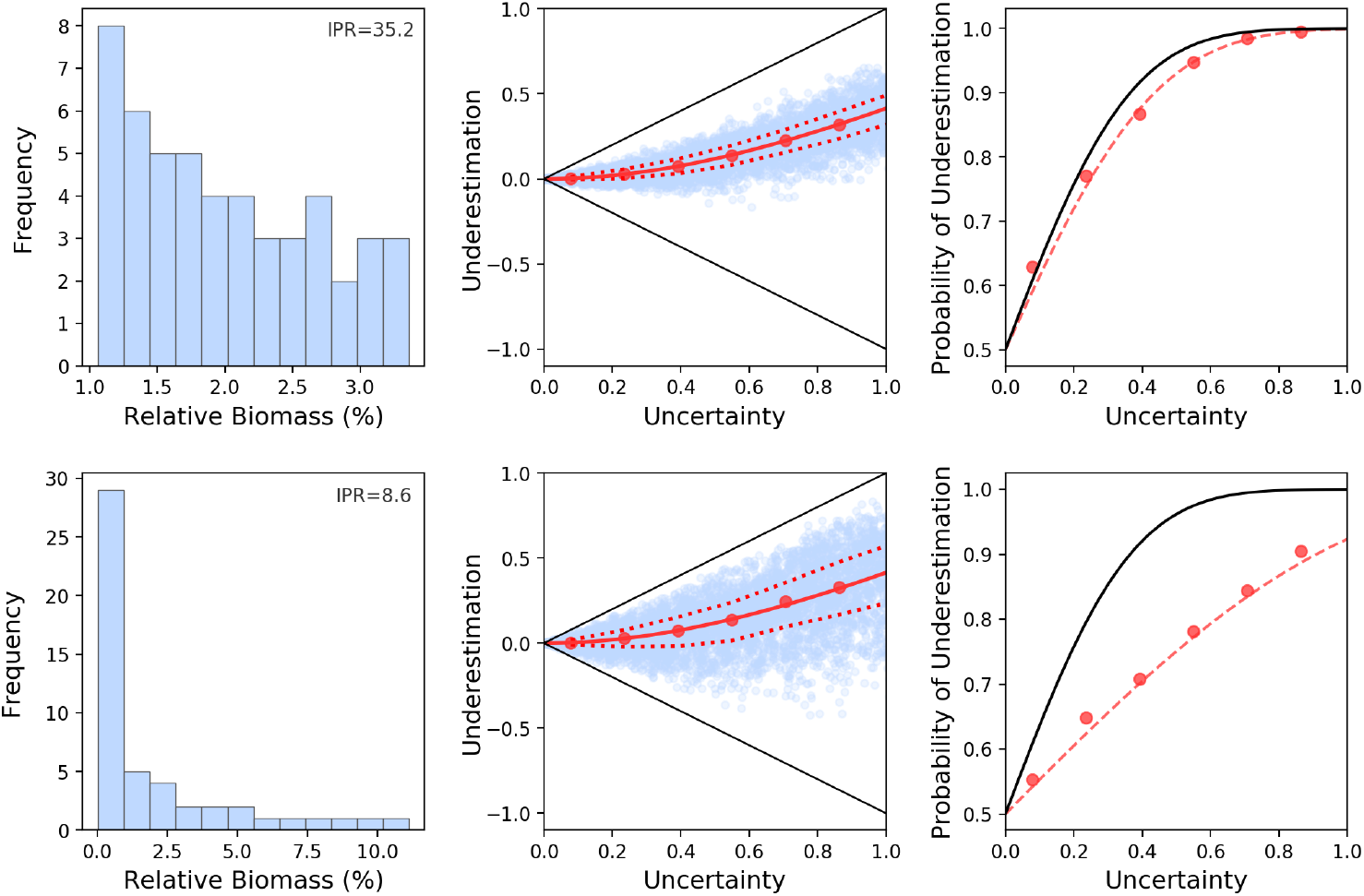
Each row corresponds to simulations of 50 species communities with uneven biomass distributions that have experienced perturbations. The first column shows the biomass distributions of these communities. The two communities have IPR, and therefore effective dimensionality, of 35.2 and 8.6. The second column shows the relationship between error and underestimation of these two communities when unbiased predictions of biomass change are scaled up to change in state-space distance. As the biomass distribution becomes more uneven the variability around the median underestimation increases (dashed lines are quantiles), which effectively reduces the probability that a given change was underestimated. This can be seen in the third column where predictions using the dimension of state-space (50, black curves) are outperformed by predictions using the IPR (35.2 and 8.6. red curves). Red points show the actual probabilities of underestimation for vertical subsets of the simulated data and are accurately predicted using the IPR.

## 3 Relevance to Ecology

### 3.1 Effective Dimensionality

We now assume that the axes that define state-space represent the biomass of the species that form an ecological system. These species may have very different abundances, and thus will not all contribute equally to a given change. For instance, in response to environmental perturbations, biomass of species typically change in proportion to their unperturbed values (Arnoldi et al., 2018; Lande et al., 2003). The more abundant species (in the sense of higher biomass) will thus likely contribute more to the ecosystem-level change. Thus, if we use species richness as a measure of dimensionality, as the above section would suggest, we will surely exaggerate the importance of rare (i.e low biomass) species. But using Eq. (5) to *define dimensionality*, we can resolve that issue. In doing so we show that the relevant dimension when applying our ideas to ecological problems is really a measure of diversity of the community prior to the change, which may not be an integer, and will typically be smaller than the number of individual components.

In fact (Appendix S3), if a species contribution to change is statistically proportional to its biomass *B_i_* the effective dimensionality of a system is the Inverse Participation Ratio (IPR) of the biomass distribution^2^, which reads:

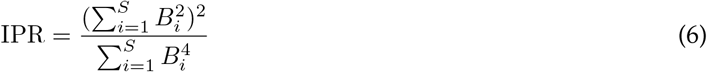

This non-integer diversity metric was developed in quantum mechanics to study localisation of electronic states (Wegner, 1980). The IPR approaches 1 when a single species is much more abundant than the others, and approaches *S* when species have similar abundance – see Suweis *et al*. (2015) where this metric is used in an ecological context. Note that the IPR is closely related (but not equivalent) to Hill’s evenness measure 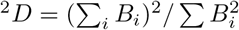 (see Appendix S3).

We can show that it is indeed the IPR that determines the variance (over a sampling of predictions and associated uncertainties of species biomasses) of the term cos*θ* in Eq. (3) so that:

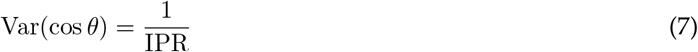

An uneven biomass distribution thus increases the width of the distribution of underestimation y therefore reducing the probability of a given realisation of change to have been underestimated. If species richness accurately predicted the width of the distribution of underestimation and therefore the probability of underestimation, the two simulated communities in Fig. 3 would behave in the same way. However, the probability of underestimation is lower than expected based on richness, particularly for the community with a more uneven biomass distribution. Indeed, replacing richness *S* by the IPR in Eq. (1) provides an excellent approximation of the behaviour of the probability of underestimation (Fig. 3).

### 3.2 Aggregate Properties and Non-Linearity

When scaling up predictions, there are different ways of measuring system-level change (Box 1). The classic reductionist approach is to quantify change via the Euclidean distance in state-space, thus keeping track of the motion of joint configurations. This is what we have done so far. Ecologically, this could correspond to measuring the absolute biomass change of a community. Here, by construction, our theory is directly relevant.

But other, non reductionist, ways of quantifying change at the system-level are possible. In ecology, this could correspond to measuring changes in the diversity, stability or functioning of the ecosystem. Yet, if differences in these properties between two states correlate with the distance in the reductionist state-space, then our theory will remain relevant. As can be seen in Fig. 4 this can be the case for diversity (Shannon’s index) and stability (invariability of total biomass (Haegeman et al., 2016)). Our theory thus applies to those ecosystem-level properties. This leads us to the conclusion that their degree of change will be systematically underestimated by predictions built from species-level predictions.

**Figure 4.**
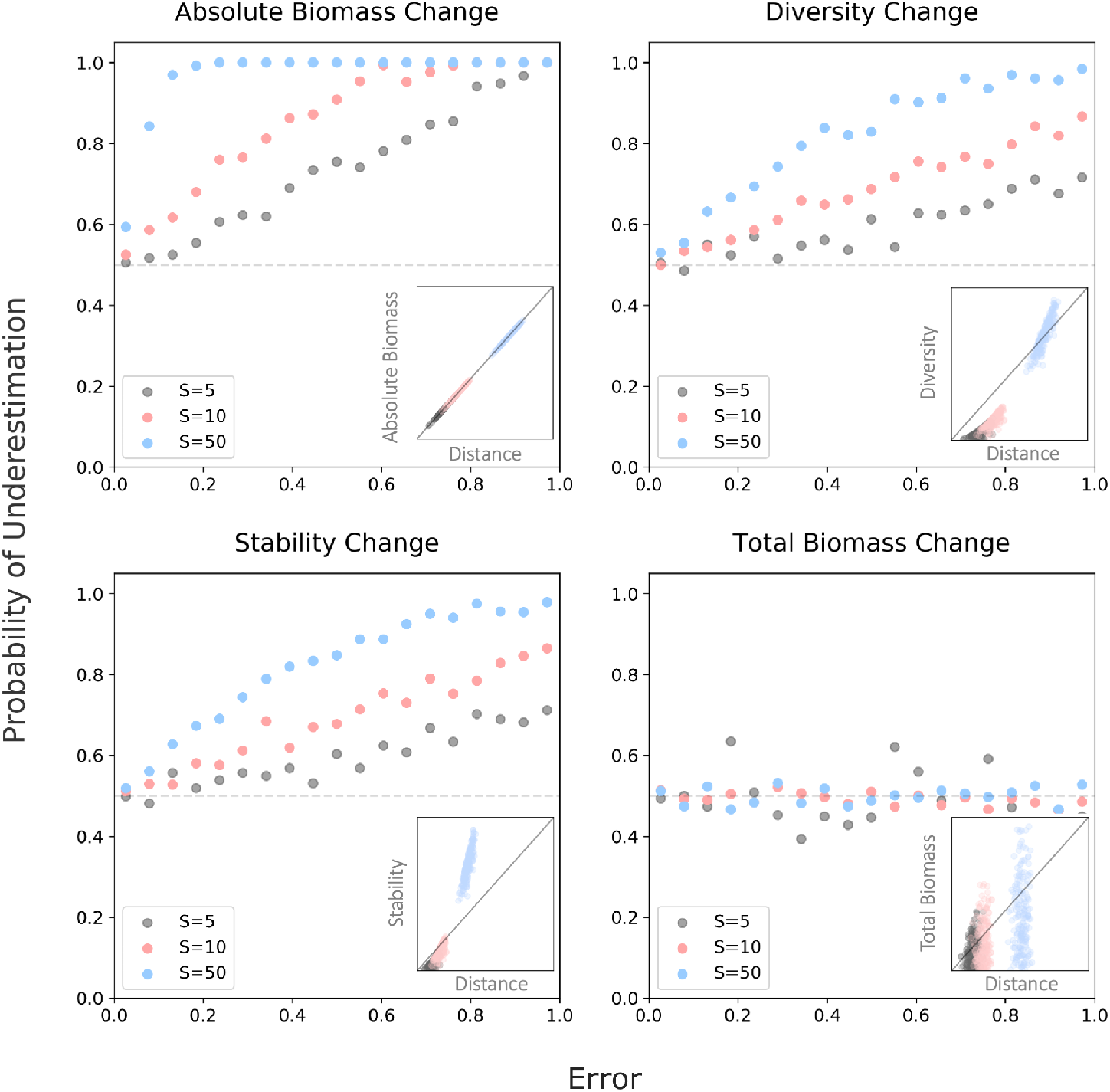
Simulated communities of 5 (grey), 10 (red) and 50 (blue) species experienced some change in their biomass. Unbiased predictions of species’ biomass change were scaled up to predictions of change in aggregate properties commonly used in ecological research. The relationship between uncertainty and the probability of underestimation is show for changes in: (1) absolute biomass, (2) diversity, specifically the Shannon index, (3) stability, specifically invariability and (4) total biomass. Subplots show the relationship between changes in each aggregate property and changes in Euclidean distance. Absolute biomass change is analogous to Euclidean distance. Diversity and stability (non-linear functions) show some correlation with Euclidean distance and are therefore sensitive to dimensional effects. Total biomass (linear function) does not correlate with Euclidean distance so scaled up predictions of change of this aggregate property remain unbiased.

On the other hand, changes of total biomass (ecosystem functioning) do not correlate well with changes in state-space Euclidean distance. This is due to the fact that total biomass is a linear function of species biomass (i.e. the sum). In fact, quantifying system-level change via a linear function acts as a projection from the state space onto a one-dimensional space defined by the function. Thus, despite the fact that the ecosystem might be constituted of many species (intrinsically high dimensional) the problem of scaling up predictions is essentially one dimensional. This is why bottom-up predictions of change of total biomass show no additional bias towards underestimation.

More generally, when the linear part of the aggregate property of interest is dominant, dimensional effects are obscured. However, as soon as we consider changes of multiple properties at once, as in multi-functionality approaches in ecology (Box 1), dimensional effects will play out – even if all aggregate properties are essentially linear.

### 3.3 Multi-Functionality

Scaling up predictions from individual components to an aggregate property can lead to a bias towards underestimation, due to dimensional effects. We explained that this occurs for non-linear aggregate properties, and not linear ones (such as total biomass). Is this to say that our theory is only relevant when predicting the change of non-linear system-level properties? Yes, but only in the restricted realm of one-dimensional approaches to complex systems.

There is, in ecology, a growing interest in multi-functionality approaches (Manning et al., 2018). These approaches are multivariate descriptions of ecosystems, an alternative to the reductionist perspective to account for the multidimensional nature of ecological systems (Box 1). By considering the change of multiple functions at once, even if these functions are essentially linear, dimensional effects will resurface.

To be clear, we still assume that we scale up predictions from the species to the ecosystem level. Only now we scale up predictions from species to several system-level properties at once, that describe the ecosystem’s state from a multi-functional point of view (Box 1). Let us suppose, for simplicity, that those aggregate properties (or functions) are essentially linear. We have seen that considering a single linear function, in terms of upscaling of predictions, essentially reduces the problem to a single dimension. Likewise, considering multiple linear functions essentially reduces the effective dimensionality to the number of functions. Subtleties arise when the number of functions (*S_f_*) and the dimensionality of the underlying system (e.g. IPR) are similar, and/or if the considered functions are colinear (see Appendix S3). For *S_f_* independent functions measured on a community we find that the effective dimensionality (the one that determines the probability of underestimation of change) is:

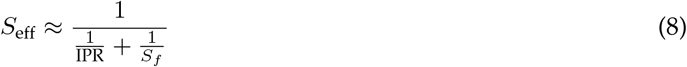

For example, if the change of an ecosystem with an IPR of 10 is measured using 10 linear functions at once, the effective dimensionality is ~5 (Fig. 5). If functions are colinear the effective dimensionality will be even lower than *S_f_*. This is to be expected, especially when thinking of an extreme case: if we measure the same function multiple times we should see no dimensional effects. In summary, in a multivariate description of complex systems, dimensional effects will inevitably play out, in more or less intricate ways, whenever a prediction is scaled up from individual components to the system-level.

**Figure 5.**
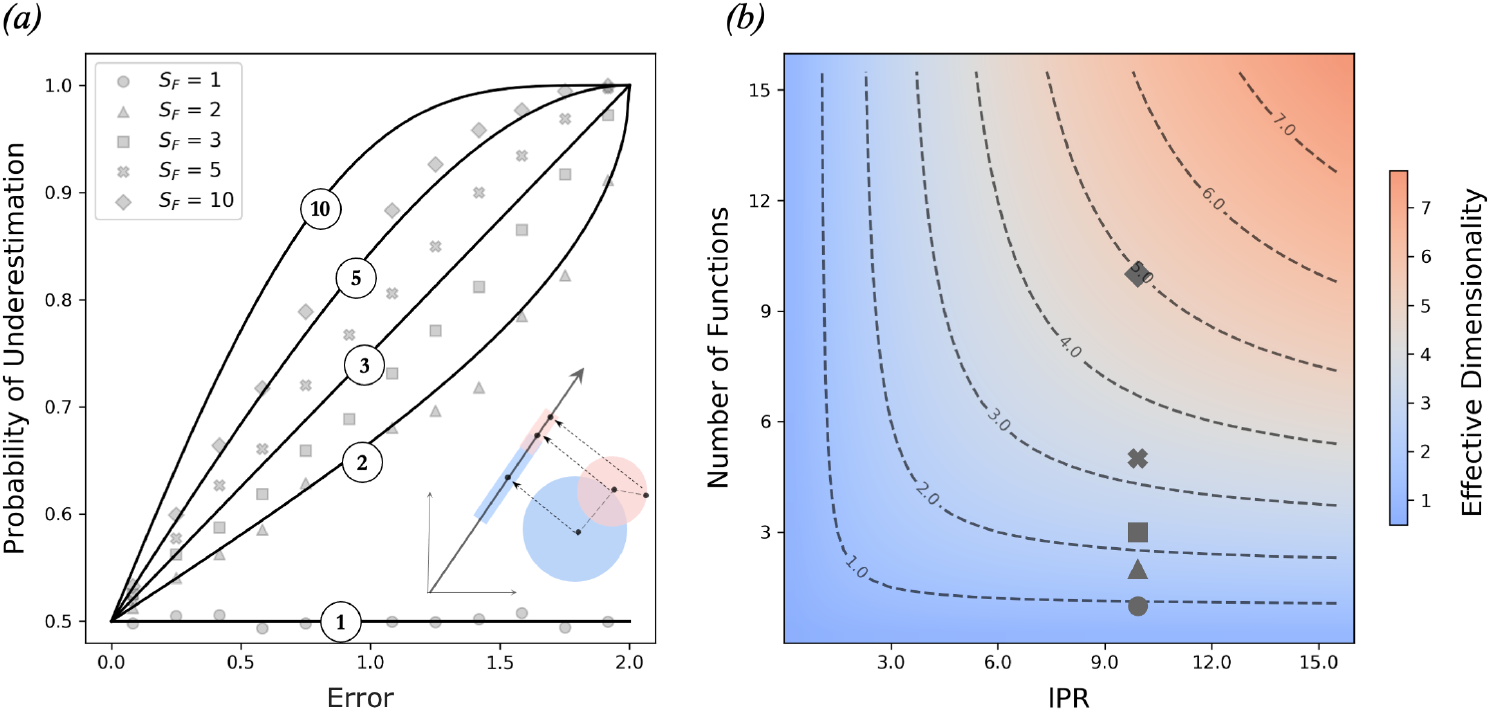
***(a)*** The relationship between prediction error caused by uncertainty and the probability of underestimation for five simulations each scaling up predictions to a different number of aggregate properties (*S_F_*). A community of 20 species, with IPR of 9.9, experienced change in biomass over 50,000 simulations. Unbiased predictions at the species level were scaled up to the community level using 1, 2, 3, 5 and 10 randomly drawn aggregate properties. Simulated results fall short of theoretical expectations for the probability of underestimation when the effective dimensionality is presumed to be the number of functions. The blue and red circles being projected onto a blue and red line represents a 2-D system being projected into 1-D functional space. ***(b)*** There is an interaction between the number of functions and the underlying dimensionality (IPR), which is illustrated by the heat-map. Usually the effective dimensionality is determined by the lower value of *S_F_* and IPR. However, when these values are similar (e.g. diamond: 10 functions and IPR of 9.9) the effective dimensionality (~5) is much lower than either value.

## 4 Discussion

Our work demonstrates that a bias towards underestimation of change emerges when predictions of individual components (e.g. species biomass) are scaled up to the system-level (e.g. ecosystem function). Our geometric approach reveals a direct relationship between the probability of underestimation, the magnitude of error caused by uncertainty and a system’s effective dimensionality. We noted that the effective dimensionality is not necessarily the number of individual components that form a system, but rather a measure of diversity *sensu* Hill (1973). In essence, these results come from the fact that *in high dimensions there are more ways to be more different, than ways to be more similar* (Fig. 6). Our goal was to make this remark quantitative and generally relevant to ecological problems.

**Figure 6.**
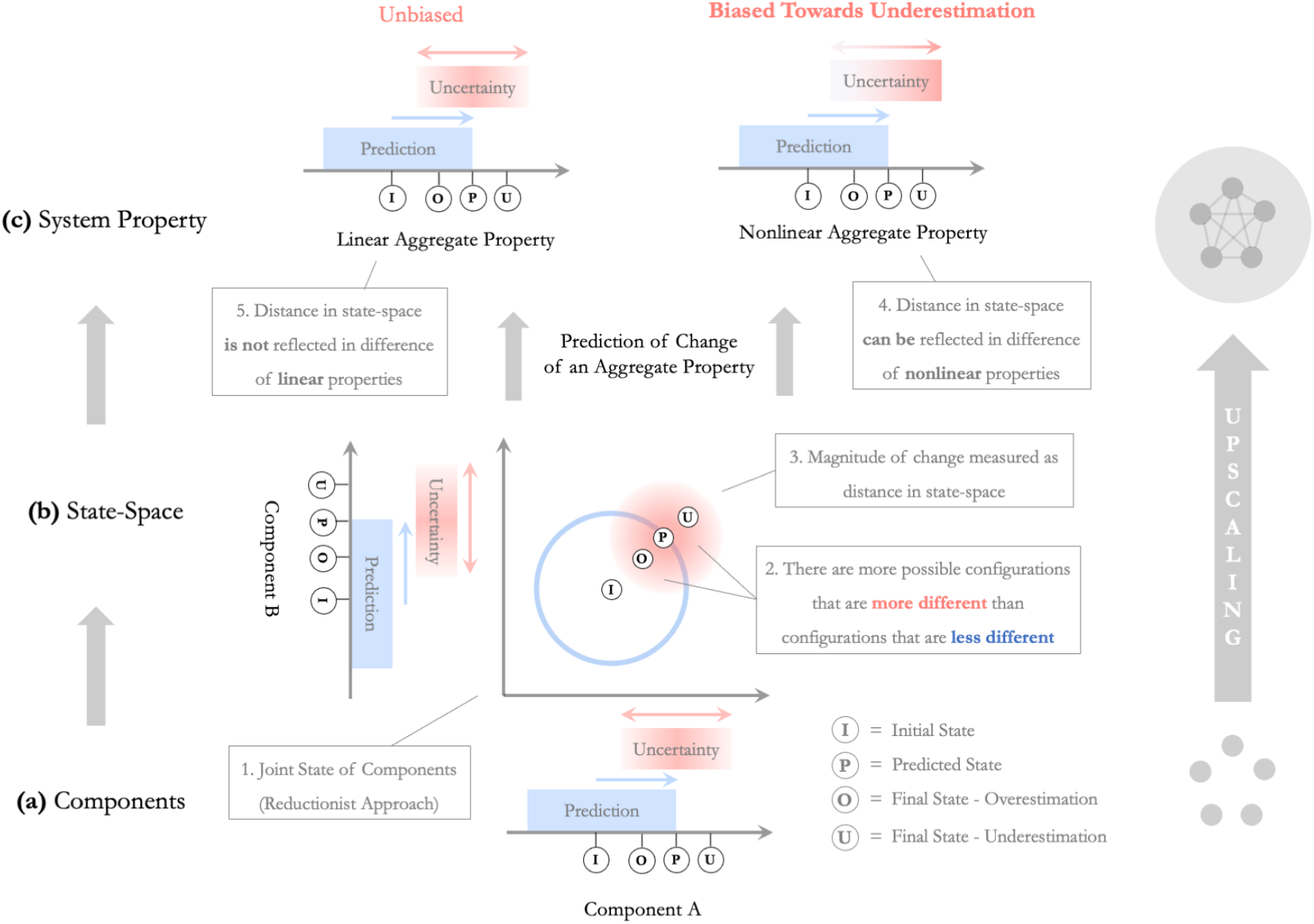
An overview of our main findings. **(a)** Two components, A and B are **(b)** considered at once to define a joint state (I). Suppose this state changes and falls near a predicted state (P). Then there are more ways for this state to be more different from (I), than ways to be more similar; more of the red disk is outside the blue circle than inside. Consequently, when predictions of change (blue) for individual components are scaled up to predictions of change of their joint state, unbiased uncertainties (red) become biased towards underestimation. In section *Geometric Approach* we quantified these surprising dimensional effects and investigate beyond the basic two-dimensional case shown here. **(c)** Magnitude of system-level change can be measured as distance in state space or by some other aggregate property. If an aggregate property is sensitive to changes in distance of the underlying state-space, dimensional effects, and therefore a bias towards underestimation, will be conserved. As we explained in section *Aggregate Properties and Non-Linearity*, it is the non-linear part of an aggregate property that controls its sensitivity to changes in state-space distance and thus the tendency of its degree of change to be underestimated by upscaled predictions.

We explained why it is non-linear aggregate properties (e.g. absolute biomass change, stability or diversity) that are sensitive to dimensional effects (Fig. 6). For linear properties (e.g. total biomass), scaling up does not generate bias. Yet, even in this case, dimensional effects will play out if several functions are considered at once to describe the ecosystem, as in multi-functional approaches in ecology.

Natural systems are intrinsically complex and the way that we describe them is necessarily multivariate (Loreau, 2010). It is generally accepted, in ecology, that there is a need for mechanistic predictive models, built from individual components and scaled up to the ecosystem-level (Harfoot et al., 2014; Mouquet et al., 2015; Poff, 1997; Woodward et al., 2010). We have shown that dimensional effects will play out in this scaling-up, generating additional bias towards underestimation of any predicted system-level change. This is not to say that scaling up predictions is a faulty approach, rather that one must keep track of dimensional effects when doing so.

Our theory provides a generic expectation for the consequences of uncertainty when predictions are scaled up from individual components to the system as a whole. As a result, it provides a baseline, of what to expect if only dimensional effects are at play, against which we can test biological (or other) effects. To inform empirical work, it is important to recognise that there are two ways that a result can deviate from our generic expectation. Focusing on the relationship between uncertainty and underestimation of change shown in Figs. 2–3, the median can be shifted due to a systematic bias caused by interactions between component uncertainties, which are assumed independent and in our framework. Furthermore, the distribution around this median can be more than or less than expected, which indicates either wrong estimation of effective dimensionality, or a systematic effect caused by something other than geometry (e.g. skewed distributions of errors or interactions). Having a clear baseline against which to identify non-geometric effects can improve our understanding of complex systems.

We only considered two levels of organisation: the level where predictions are made and the level where predictions are scaled up to. However, intermediate levels could, in principle, be considered. For instance, given the increasing resolution of ecological data, predictions of change may originally be based at the level of individual organisms and could first be scaled up to species-level predictions and subsequently scaled up to ecosystem-level predictions. Here, if non-linear aggregate properties are used, dimensional effects will bias species-level predictions towards underestimation and will further increase this bias for ecosystem-level predictions. With an ever-increasing resolution of data, scaling predictions across multiple levels of organisation, and potentially introducing dimensional effects at multiple levels, may become more common in the study of complex systems.

Our work is theoretical and, in essence abstract. Yet it may be relevant for highly practical domains of ecology. To make this point, we will now discuss some implications of our theory to multiple-stressor ecological research, an essentially empirical field that explicitly deals with considerable uncertainty of predictions and holds great interest in its consequences.

### 4.1 Multiple-Stressor Research

In the light of our theory, we propose to revisit a seemingly unrelated problem of wide ecological interest: what is the combined effect of multiple stressors on a given ecosystem? By translating our theory into the language of multiple-stressor research we aim to highlight some implications and to inspire further generalization.

The combined effect of stressors on an ecological system is generally predicted based on the sum of their isolated effects, i.e. an “additive null model” (Folt et al., 1999; Schäfer and Piggott, 2018). Uncertainty around this additive prediction, which is ubiquitous in empirical studies (Crain et al., 2008; Holmstrup et al., 2010; Jackson et al., 2016), causes prediction errors called “non-additivity”. Uncertain predictions will either overestimate or underestimate the combined effect of stressors, respectively creating “antagonism” and “synergism” (Folt et al., 1999; Piggott et al., 2015). This translation of stressor interactions in terms of prediction uncertainty and under- or over-estimation lead us to the conclusion that scaling up uncertain multiple-stressor predictions generates bias towards synergism.

Here, scaling up predictions refers to multiple-stressor predictions (e.g. an additive model) at one level (e.g., individuals, populations) being used to build multiple-stressor predictions at higher levels of biological organisation (e.g. communities, ecosystems), an approach for which there is growing interest (Côté et al., 2016; Kroeker et al., 2017; Orr, Vinebrooke, et al., 2020; Thompson et al., 2018). To be clear, scaling up predictions is not equivalent to simply scaling up investigations; our theory does not predict greater synergism at higher levels of organisation. In fact, we are not making predictions about how stressors will behave at higher levels of organization. What we claim instead is that, if we have a model for the combined effect of stressors at one level of organization and use that model to deduce their combined effect at higher levels, the process of scaling up the model will introduce a bias towards an observed synergy between stressors, even if no systematic synergy was observed at the lower level.

Our theory has consequences for the interpretation of stressor interactions and is therefore relevant to the debate surrounding multiple-stressor null models (De Laender, 2018; Griffen et al., 2016; Liess et al., 2016; Schäfer and Piggott, 2018). Our findings are especially relevant to the *Compositional Null Model*, which employs a reductionist approach to the construction of multiple-stressor predictions (Thompson et al., 2018). In such an approach, the baseline against which biological effects are tested must be shifted. Dimensional effects, quantified by the effective dimensionality of the underlying system and the non-linearity of aggregate properties, need to be accounted for to decipher a biological synergism from merely a statistical synergism.

### 4.2 Conclusions

In this paper we have addressed a subproblem of the reductionist program (Levins and Lewontin, 1985; Loreau, 2010; Wan, 2013). We investigated the consequences of uncertainty when unbiased predictions of individual components are scaled up to predictions of system-level change. Due to a geometric observation that *in high dimensions there are more ways to be more different, than ways to be more similar*, scaling up uncertain predictions can underestimate system-level change. These dimensional effects manifest when non-linear, but not linear, aggregate properties are used to measured change at the system level, and when multiple functions are considered at once. Although we have primarily focused on ecology, and in particular on the response of ecosystems to perturbations; our general findings could inform any field of science where predictions about whole systems are constructed from joint predictions on their individual components, such as economics, finance, energy supply, and demography (Box 2).

#### Box 2: Generalisation Beyond Ecology

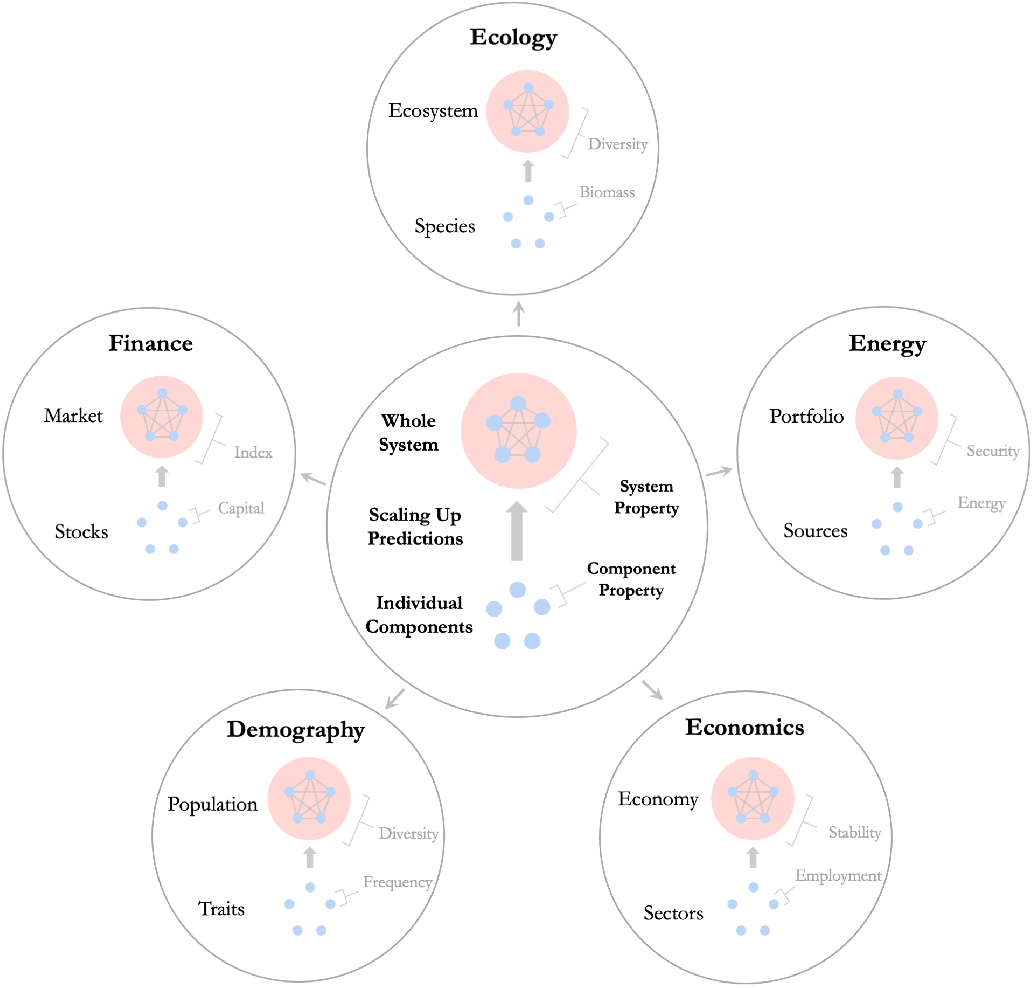

Our findings could be relevant to other fields of science where: (i) there is interest in predicting change of complex systems based on knowledge about their individual components, and (ii) systems are described using multivariate coordinates and/or using non-linear properties of individual components.

- In **economics**, a region’s economy can be viewed as a complex system comprised of individual sectors (e.g. agriculture, tourism, technology). Predictions of how employment numbers will change in individual sectors due to some perturbation could be scaled up to predictions of change of economy-level properties of interest such as stability, measured as, for example, the evenness of employment across sectors (Dissart, 2003; Halpern et al., 2012; Malizia and Ke, 1993).
- In the study of **energy supply**, different fuel or energy sources of a country (e.g. solar, wind, oil) can be considered together in a country’s energy portfolio. Predictions of change of energy generation in each individual source could be scaled up to predictions of change of portfolio-level properties. Energy security is a system-level property of great interest that is quantified using diversity metrics (Chalvatzis and Ioannidis, 2017; Stirling, 1994) or variance-based approaches (Roques et al., 2008) based on *Mean-Variance Portfolio Theory*, which was originally developed to study risk or volatility of investment portfolios (Markowitz and Todd, 2000).
- In **demography**, populations can be thought of as systems comprised of multiple different groups that are defined by traits (e.g. gender, age, ethnicity). Again, diversity is a system-level property of great interest in the study of populations that is quantified using non-linear aggregate functions (Reardon and Firebaugh, 2002; White, 1986). Changes in diversity of human populations is pertinent to many social sciences including **sociology, economics and politics**.
- In **finance**, markets are complex systems whose individual components are stocks. Predictions of how the capital of individual stocks will change could be scaled up to predictions of how stock market indices will change. Certain stock market indices, for example diversity-weighted indices, are non-linear aggregate properties that will be sensitive to dimensional effects (Chow et al., 2011; Fernholz et al., 1998). At a different financial scale, our theory may also be relevant to the study of investment portfolios. Here, analogous to energy security, portfolios are systems comprised of individual assets and the volatility or risk tolerance of a portfolio (measured using non-linear aggregate properties) is of great interest to investors (Bera and Park, 2008; Markowitz and Todd, 2000).

## Data accessibility

All data were simulated. Code is available in a Jupyter Notebook on GitHub: https://github.com/jamesaorr/scaling-up-uncertain-predictions.

## Supplementary material

Code is available in a Jupyter Notebook on GitHub: https://github.com/jamesaorr/scaling-up-uncertain-predictions.

## Acknowledgements

We thank Matthieu Barbier, Nuria Galiana and Yuval Zelnik for discussions and review of previous versions of this work. JFA and ALJ were supported by an Irish Research Council Laureate Award IRCLA/2017/186. JO was supported by an Irish Research Council Laureate Award IRCLA/2017/112 and TCD Provost’s PhD Award held by JP. Version 3 of this preprint has been peer-reviewed and recommended by Peer Community In Ecology (https://doi.org/10.24072/pci.ecology.100063)

## Supporting Information

### S1 Geometrical model

Consider a complex system whose states are given by points in *R^S^* (thus determined by *S* individual variables, e.g species biomass). Let *v* ∈ *R^S^* be an expectation for a change of state. Let *w* be the actual change that is observed, and define the error vector *u* such that *w* = *v* + *u*. From *u* and *v* we define a scalar measure *x* of relative error as

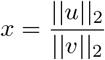

We formalize the question of whether there has been more change observed than predicted, by defining

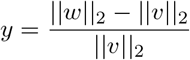

In both of the above expressions || · ||_*p*_ denotes the *L_p_* norm of vectors. *p* = 2 corresponds to Euclidean distance, we still see bellow that other values of *p* can occur in our formalism. Also, our results hold for other choices of norm in defining *x* and *y*. The Euclidian norm is however, the most convenient for a geometrical approach. A reorganization of *y* gives

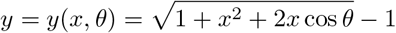

where *θ* is the angle between error *u* and prediction *v*, that is

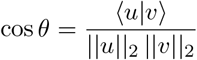

### S2 Random ensemble

We now assume that *u* and *v* are random variables (but the prediction *v* could also be given). We assume however that the components of *u_i_* have zero mean and median – the prediction of individual variables is unbiased and unskewed. Then *E_u_* 〈*u*|*v*〉 = 0, thus *Ecosθ* = 0. This implies that

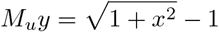

At fixed error *x*, the distribution around this median is driven by variance of cos*θ*, over random draws of vectors *u* and *v*. We first define the covariance matrices 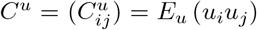, and 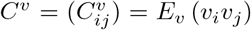. We then have that

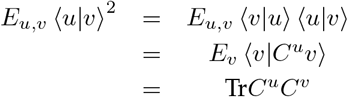

and similarly

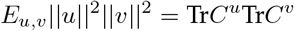

Thus

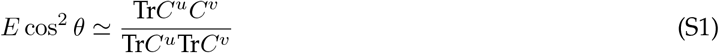

**Example**

Suppose that *C_u_* = *σ*^2^*I* where *I* is the identity matrix. This implies that uncertainties of the individual variables are independent random variables with variance *σ*^2^. We then have

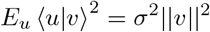

while

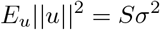

so that

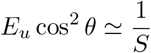

#### S2.1 Probability of underestimation

Given an imprecision level *x*, the theory has underestimated the actual response if *y*(*x, θ*) ≥ 0 and thus if the angle *θ* between the theoretical prediction *v* and the vector of unaccounted change *u* satisfies

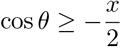

If cos*θ* is approximately normally distributed with zero mean and variance 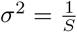, than

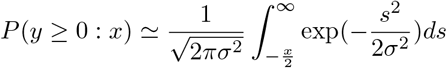

hence, by the properties of the cumulative distribution function of standard normal distributions, one gets

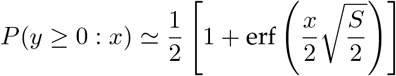

where erf is the error function. This expression should be compared to the exact solution in the case of a uniform sampling over the direction of *u* (which is the case if 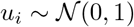 –uncertainties of individual variables are independent and normaly distributed). In this special case the problem of deriving the probability of underestimation becomes purely geometrical: it is the surface of a ball of radius *x* and centered on the unit sphere, that is contained in the unit ball. One then gets

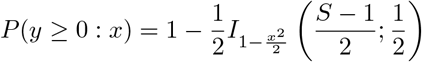

Where *I_s_*(*a, b*) is the regularized *β*–function (the cumulative distribution of the *β*-distribution). In fact those two expression converge at high diversity *S*. In any case, we see here that the probability of underestimation will grow with *S*.

### S3 Effective diversity

*S* may not always be the relevant measure of diversity. Indeed if 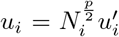 where *N_i_* is the abundance (or biomass) of species *i* and *C^u′^* ∝ *I* then *C^u^* ∝ *D^p^* where *D* is a diagonal matrix with *D_ii_* = *N_i_*. If *v_i_* obeys a similar rule, so that *C^v^* ∝ *D^q^* then

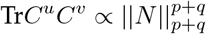

while

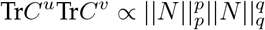

so that

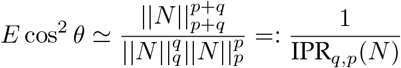

In particular, for *q* = *p* = 2 we get that

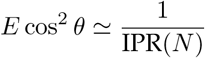

where IPR(*N*) is the Inverse Participation Ratio, a measure of diversity of the abundance distribution *N*. The more general expression above can also be seen as a measure of effective diversity. It can be compared to Hill’s diversity metrics with index *Q* = *p* + *q*

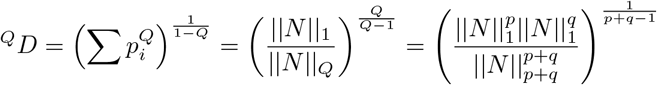

where *p_i_* is the relative abundance of species *i*. We indeed see that *^Q^D* coincides with IPR_*q,p*_ when *q* = *p* =1, and stays closely related in general. In fact, using the inequality

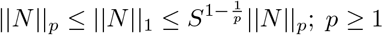

one gets, for *p*, *q* ≥ 1

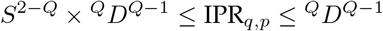

#### S3.1 Probability of underestimation

If cos*θ* is approximately normally distributed with zero mean and variance (where *S*_eff_ ≤ *S* would be an effective dimensionality as defined in the previous sections), than

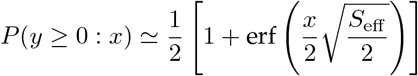

This expression should be compared to the exact solution derived above in the case of a uniform sampling over the direction of *u* (the case if 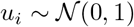), which suggest the Ansatz

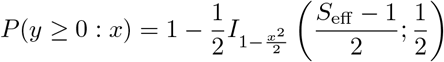

when the effective dimensionality is not necessarily *S* or even an integer (the two expressions uniformly converge towards one another as *S*_eff_ grows).

#### S3.2 Projection on linear functions

Suppose now that we measure *S_F_* linear functions of species biomass of the form

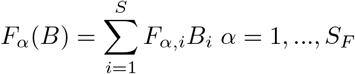

We must now project the covariance matrices onto the space spanned by the gradient (*F_α_,i*) of the functions. If *P_α_* = (*F_α,i_F_α,j_*) the projector on the function *F_α_*, we can do this as

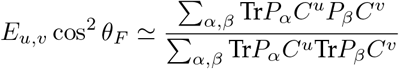

When taking a ensemble average of functions, with 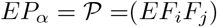, we must take care in differentiating terms in sums for which *α* ≠ *β* and terms where *α* = *β*. In the former case the projectors *P_α_* and *P_β_* are independent random variables and we can replace them by their mean 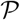. In the latter case, we must first define 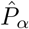 as the linear operator that maps a matrix *M* to *P_α_MP_α_*; its ensemble mean 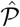 encodes the 4th moments of *F_α_*. We then get

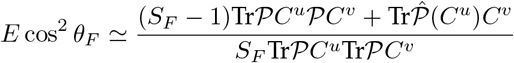

**Example 1**

This example is the one treated in the main text, where the functions are statistically independent of one another. Suppose as before that *C^u^* = *C^v^* = *D*^2^ (*p* = *q* =1) and *EF_i_* = 0 *EF_i_F_j_* = *δ_ij_* (this condition is what we mean by statistically independent). One gets that

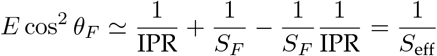

so that at first order, the effective dimensionality *S*_eff_ is the harmonic mean

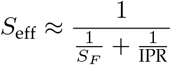

as presented in Eq. (8)

**Example 2**

For the sake of completeness we treat here the case where the functions are not statistically independent due to the fact that *m*_1_ = *E*(*F_j_*) ≠ 0 (the average species contributions to functions tends to be either systematically positive or negative). In this case 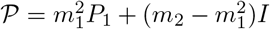, where *P*_1_ is a matrix whose elements are all equal to 1, and *m_n_* are the *n*-th moments of *F_i_*. We have that

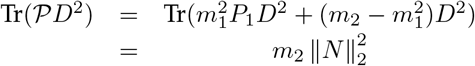

and so

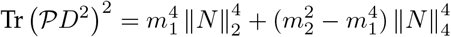

on the other hand, one can show that

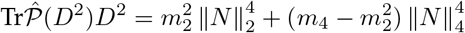

if

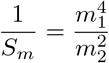

we get that

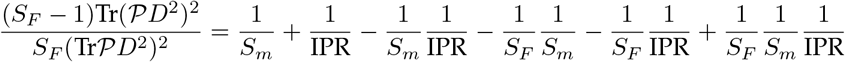

on the other hand

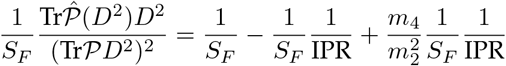

summing the two gives

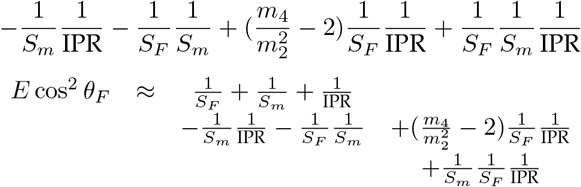

for a normal distribution

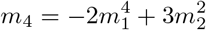

thus

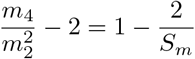

we then have

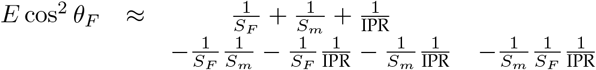

We see here interactions between the various dimensions *S_m_*, *S_F_* and IPR, with a potential dominance of *S_m_* when all other are much larger. This effective dimensionality emerges due to the collinearity of functions, which thus span a subspace of potentially much smaller dimension than *S_F_*.

#### S3.3 Change of metric

Consider a non euclidean metric tensor *H* (i.e a positive definite matrix). Distances must now be measured as

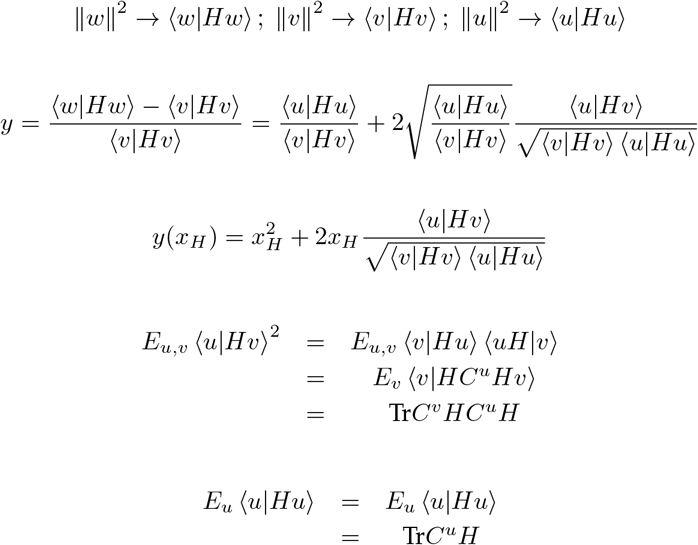

Thus

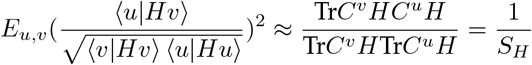

the change of metric can thus change the effective dimensionality. In particular, if *C^u^* ∝ *C^v^* ∝ *I* this gives

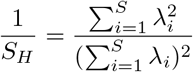

where λ_i_ are the eigenvalues of *H*. Note that *H* could be the Hessian function (second derivatives) of a non linear function, computed near the initial state. This explains how non linear functions can induce a dimensionality effect on the probability of underestimating change, as illustrated in Fig. S1.

**Figure S1.**
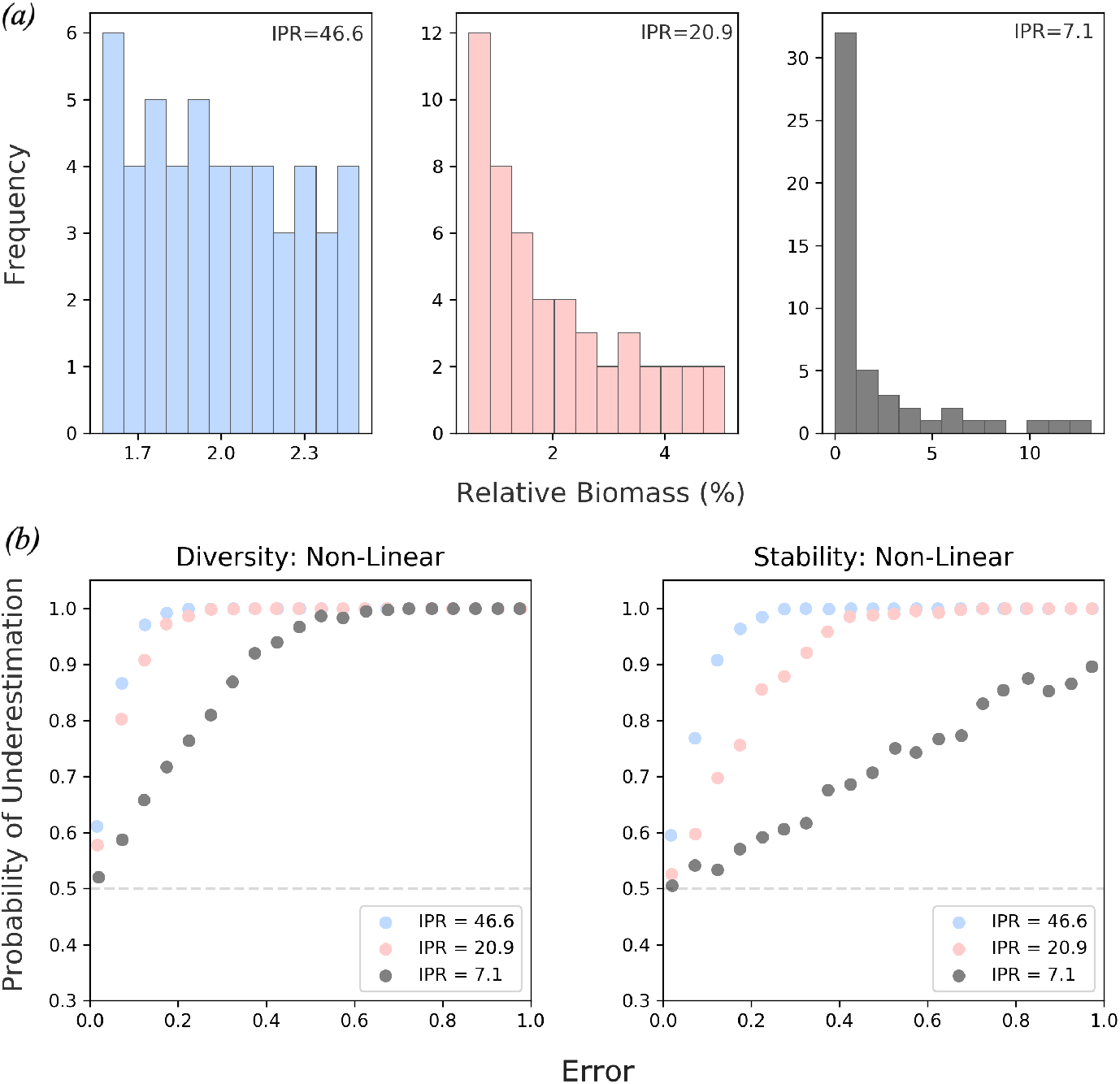
***(a)*** Biomass distributions of three 50-species communities with IPR and therefore effective dimensionality of 46.6 (blue), 20.9 (red) and 7.1 (grey). **(b)** The non-linear contribution of diversity (the Shannon index) and stability (invariability) towards the probability of underestimation; it is the non-linear part of a function that is sensitive to the dimensionality of the underlying system.

### S4 Simulations

Initially, the theoretical relationship between error, underestimation and dimensionality was tested using numerical simulations (Fib. 2(c)). These simulations uniformly sampled the intersecting circles, spheres and hyper-spheres defined by a prediction of change and relative error (Fig. 2). This was done for 1-D, 2-D, 10-D and 20-D systems over 20,000 simulations. Specifically:

- a prediction of change and was randomly generated from a normal distribution of mean 0 and standard deviation 1 (defining the blue circle in Fig. 2a).
- a direction of error was randomly generated from a normal distribution of mean 0 and standard deviation 1, and a magnitude of error was randomly generated from a uniform distribution between 0 and 2 (defining the the red circle in Fig. 2a).
- From these values, error (*x*) and underestimation (*y*) were calculated based on Euclidean distance and subsequently plotted in Fig. 2c).
- The probability of underestimation *P*(*y* > 0; *x*) was calculated from the simulated results of error and underestimation.

As a next step, these simulations were modified to fit ecological problems. In Fig. 1 and Fig. 4 the intersecting shapes that are uniformly sampled had dimensions determined by the number of species in a simulated community. However, the dimensions of state space were given unequal weighting of how they respond to change in the form of uneven biomass distributions randomly generated from a log normal distribution of mean 0 and standard deviation 0.05.

In Fig. 3 and Fig. S1 communities of 50 species were given unequal biomass distributions by drawing species’ biomass from a log scale of varying range; the wider the range of the log scale the more uneven the biomass distribution. Underestimation (y) was calculated using Euclidean distance *and* a number of ecological relevant aggregate properties: the Shannon index (diversity), invariability (stability) and total biomass (functioning).

For Fig. 5 our simulations were modified to illustrate that additional dimensional effects come into play when changes in multiple functions are considered at once. Over 50,000 simulations 20-D hyper-spheres (community of 20 species) with unequal weighting (IPR of 9.9) were uniformly sampled and the results were projected into functional space. Specifically, underestimation was measured for 1, 2, 3, 5 and 10 aggregate functions. Linear aggregate functions of the form:

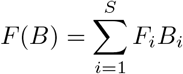

were defined via the coefficients *F_i_*, i.e. their sensitivity to the change in the biomass of species *i*. The sensitivity of an aggregate function to each species was randomly drawn from a normal distribution of mean 0 and standard deviation 1. This corresponds to the case of statistically independent functions (see example 1 in subsection S3.2). State space was then defined by the number of functions.

Simulations were conducted in Python with the Matplotlib, NumPy and SciPy libraries. Code is available in a Jupyter Notebook on GitHub: https://github.com/jamesaorr/scaling-up-uncertain-predictions.

## Footnotes

1 This is the most convenient norm for our geometrical approach but other norms would give similar results.

2 our theory allows other choices of statistical relationships between biomass and contribution to change, leading to other diversity metrics, which can be seen as generalization of the Inverse Participation Ratio.

